# Multifractal signatures in ossification reveal a disordered-to-ordered transition in mineral patterns

**DOI:** 10.1101/2023.08.31.555718

**Authors:** Mohammadreza Bahadorian, Johanna Lattner, Jacqueline M. Tabler, Carl D. Modes

**Affiliations:** Max Planck Institut for Molecular Cell Biology and Genetics (MPI-CBG), 01307 Dresden, Germany; Center for Systems Biology Dresden (CSBD), 01307 Dresden, Germany; Cluster of Excellence, Physics of Life (POL), TU Dresden, 01307 Dresden, Germany

## Abstract

Developing biological systems can exhibit both dynamic pattern formation and cross-scale interactions. Multiscale relationships are critical in the establishment of these patterns but remain poorly understood. Classification of mineral pattern in bone is a quintessential example. One approach to quantifying these patterns relies upon statistical self-similarity and, in particular, monofractal analysis. However, simple monofractal characterisations fail to capture the complexity of multiscale interactions in developing biological systems. Here we show that multifractal techniques, effectively capture the complex patterns of self-similarity in a dimensionally reduced, usable way. Further, we show that a simple generative model of ossification in the mouse skull, coupled with multifractal methods indicates a primary role of collagen density in pattern establishment and predicts the existence of a sharp boundary in pattern complexity.

## I INTRODUCTION

Patterns across biology frequently confer function. Despite the familiarity of these patterns, there remains little understanding of how they can be generated across scales. Although classical Turing patterns and broader reaction-diffusion systems provide a well-explored example of pattern formation that spans scales [1–3], the establishment of many multiscale patterns in biology cannot be explained through this class of mechanism. In particular, a hallmark of mesoscale patterns (e.g. as in organs) is not only their emergence from dynamics at smaller scales but also their significant structural complexity (e.g. as in organs).

One might hope that the pattern of morphogenesis at the mesoscale could be leveraged to understand how smaller-scale interactions combine to drive emergence. However, there is so little known about these dynamics that different approaches to exploring pattern generation and establishment are required. Indeed, characterization of patterns themselves is in many cases highly non-trivial and in need of better, more unified approaches for quantification and decomposition. This kind of improvement in pattern analysis holds great potential, as in fluid flow in patterning vasculature, where analyzing patterns unveiled the primary role of wall shear stress in vascular morphogenesis [4–6].

One quintessential organ system that emerges from multiscale interactions and whose function emerges from its pattern is bone [7]. Collagen, 1a1, the major building block of bone, mediates these cross-scale interactions. The nanoscale structure of collagen laid down by osteoblasts determines the coarse-grained material properties of skeletal elements [8, 9]. Mineralization then canalizes these multiscale interactions as carbonate-substituted hydroxyapatite crystallizes on collagen fibrils to generate the bone’s ultimate form and mechanical characteristics [10–12]. Could collagen, therefore, be the key to understanding the emergence of complex bone pattern? Unfortunately, here one encounters a signficant obstacle to understanding the formation of flat bones: the enormous variability and spatial fluctuations arising from the random timing of cellular events (*e*.*g*. differentiation, motion, and division) across the tissue throughout development [13, 14].

If the nanoscale properties of collagen indeed give rise to mesoscale bone patterning during morphogenesis, cross-scale analyses would be required to understand ossification. However, converting mesoscale pattern description into mechanistic understanding on the smaller scales is a major challenge without decomposition or reduction of complexity [15]. The traditional bottom-up approaches of reduced, predictive, theoretical models employed to understand emergence from few stereotypic interactions fail without appropriate simple metrics with which to compare model output to complex data. At the same time, higher-order analyses of complex data are highly non-trivial and, in many cases, the classical approaches such as fractal scaling, cannot reflect the symmetries and character of the data, rendering them ineffective. For this reason monofractal approaches are not currently used for diagnosis of bone diseases despite their success at the research stage [16, 17]. Moreover, while monofractal approaches allow for a single multiplicative scale of self-similarity, multifractal approaches, on the other hand, can take into account a spectrum of such simultaneous scaling behaviors [18]. Therefore, multifractal analyses may present a way forward, as they are capable of capturing higher-order and multiscale features of complex data in a general way [19]. Further, multifractal analyses can also provide the simple metrics needed for reduced models to be effective.

Here, with a simple generative model we reproduce a segment of ossification patterns of the developing mouse skull vault. We then show the utility of two multi-fractal analysis methods (Multi-Fractal Detrended Fluctuation Analysis, MFDFA and Wavelet Transform Modulus Maxima Maxima Method, WTMMMM) to describe mineralisation patterns in mouse embryos as well as the simulation results. Using MFDFA, we find emergent mineral pattern indeed has multifractal features. This multifractality can in turn be used to characterize emergent patterns. We also investigate the origin of the observed multifractality by employing data surrogation techniques. By studying our simulation results via MFDFA we predict that collagen density in particular is a key regulator of mineral pattern while its organization is not. We experimentally confirm this hypothesis by demonstrating the sufficiency of nanoscale collagen density in generating multifractal mineral patterns during development. Furthermore, our generative model predicts the existence of a transition boundary from multifractality to monofractality in ossification patterns.

## II RESULTS

Expansion of mineralized frontal and parietal bones from the lateral aspect of the forebrain to the medial apex of the head occurs during the last half of embryonic development (Fig. 1A). In the mouse, embryonic day (E) 14.5 marks the first appearance of mineral that can be stained with Alizarin red and which forms a sparse lacelike pattern (Fig. 1B, C). Over the following 5 days, this mineral expands towards the midline, and spaces within the growing meshwork progressively fill to generate new pattern as the bone matures (Fig. 1B, D). While developmental studies commonly leverage this pattern in phenotypic analyses and find qualitative differences in perturbation schemes [], no quantitative tools have been implemented to visualize and describe these patterns over time. Quantifying mineral pattern in whole embryos is challenging due to curvature of the cranial bone pairs over the brain, therefore, we excised stained skull caps and captured two-dimensional images of frontal and parietal bone pairs in flatmount. This allowed us to image the autofluorescence of Alizarin red stained mineral at high-resolution to generate monochrome images (Fig. 1B-D) for downstream computational analyses (Fig. 2A step 1-5).

**FIG. 1:**
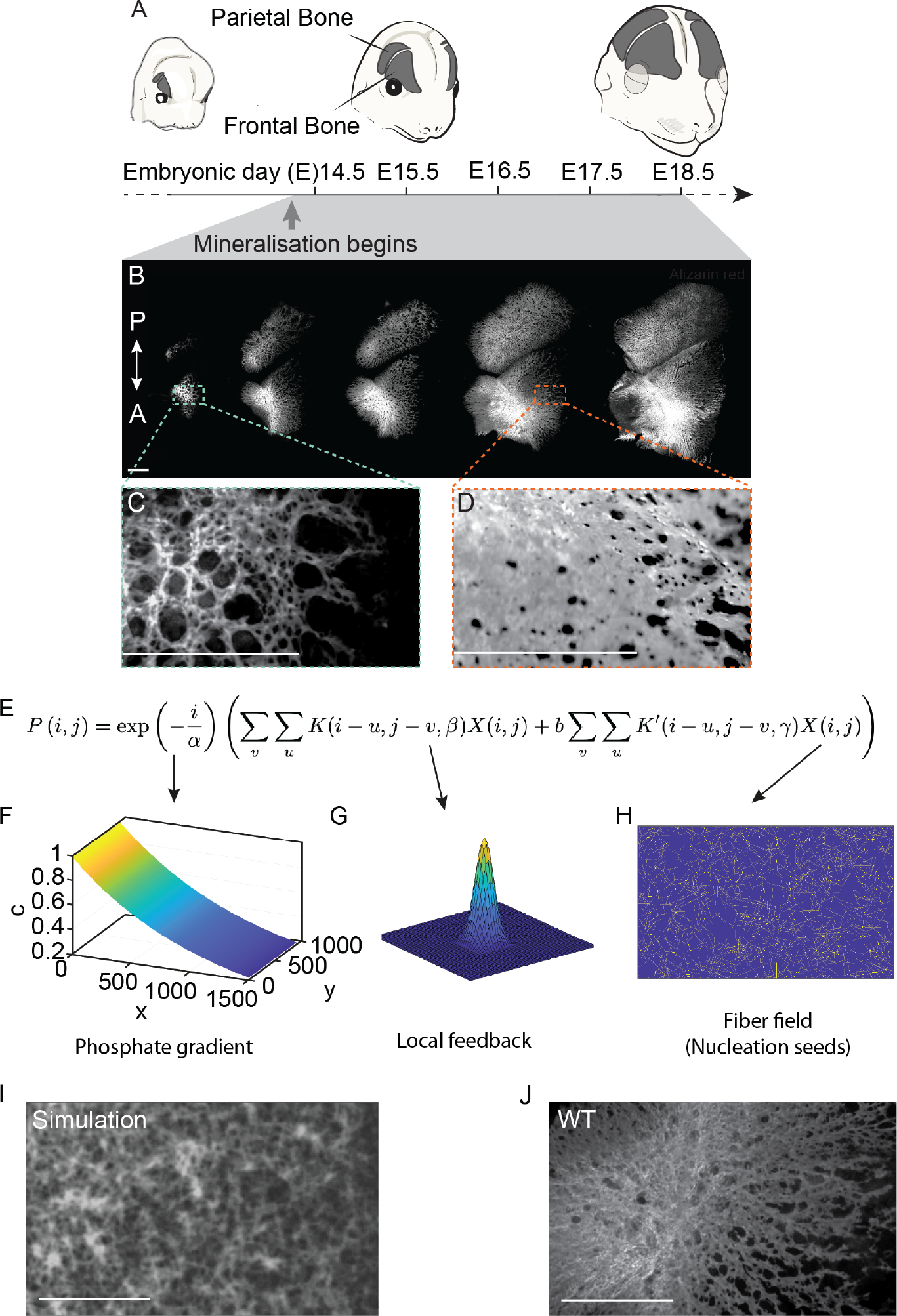
Ossification patterns throughout development and the proposed generative model. **(A)** Schematic illustrating frontal bone expansion toward the midline during morphogenesis of the mouse skull vault which begins at E13.5 and continues until E17.5 when frontal bones meet at the midline joints (termed “sutures”). Mineralisation begins at E14.5. **(B)** Alizarin red staining of mineral in excised whole-mount frontal bones at E14.5, E15.5, E16.5, E17.5, and E18.5, respectively from left to right. (A, anterior. P, posterior). The dashed boxes denote insets presented in **(C)** and **(D)**. Magnified views of E14.5 and E17.5 Alizarin red staining, respectively. Scale bars, 500*µ*m. **(E)** The probability of ossification in each iteration at lattice site (*i, j*). **(F)** One-dimensional exponential pre-factor captures the effect of the phosphate gradient. **(G)** Positive local ossification feedback term, mediated by a Gaussian kernel. **(H)** Random collagen fiber field nucleation template term. **(I)** Typical simulation results. **(J)** A segment of a typical WT E15.5.

**FIG. 2:**
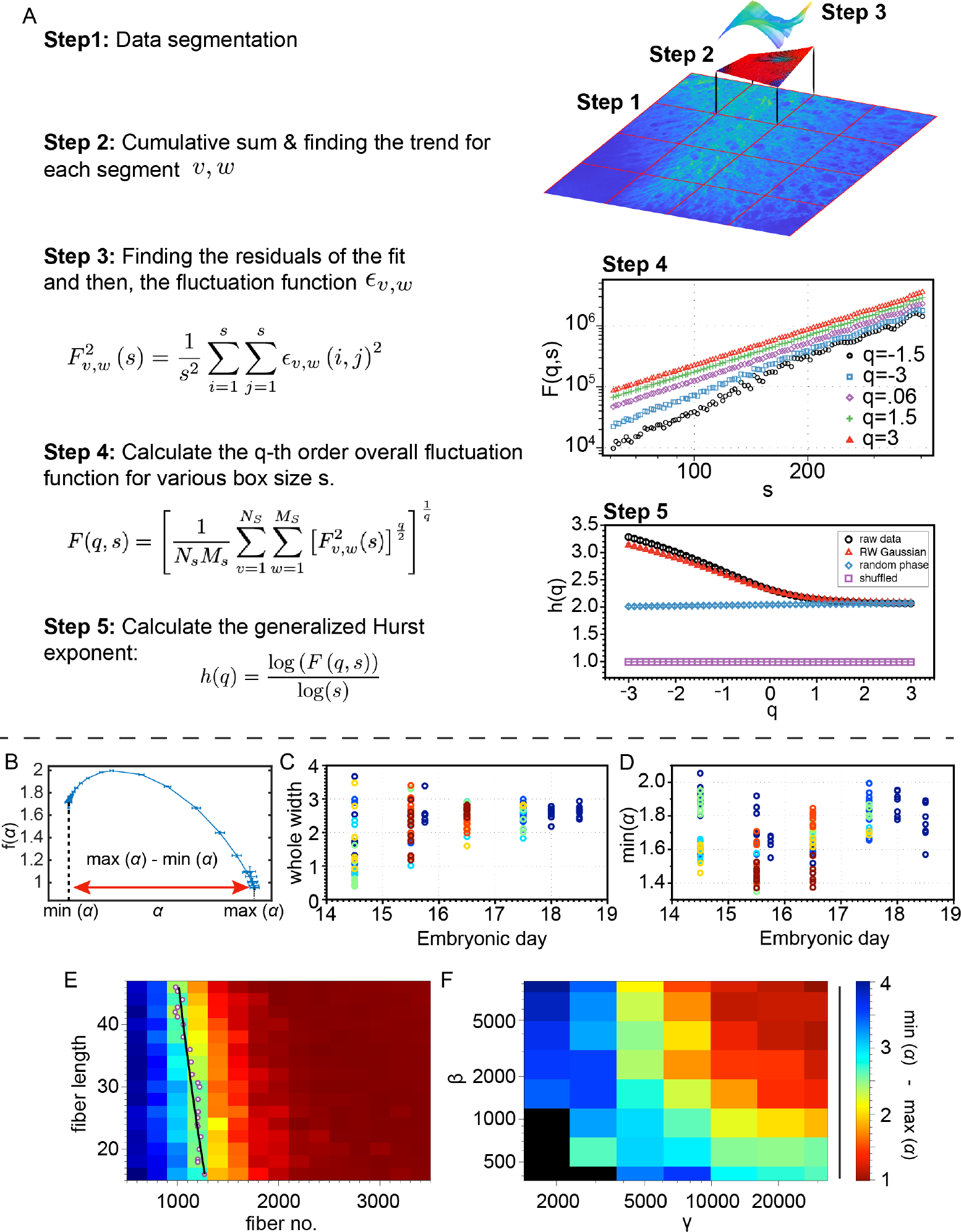
Singularities and singularity detection in data. **(A)** schematic steps of MF-DFA. **(B)** A typical singularity spectrum and the definition of its characteristics.**(C)** Multifractality defined as Δ*α* := max(*α*)*−* min(*α*) over development. **(D)** The most irregular singularities of the ossification patterns over development. **(E)** Multifractality vs. density of collagen fibers. **(F)** Multifractality vs. length scale of the exponential determining phosphate gradient *β* and length scale of the exponential determining fiber density *γ*.

Despite the complexity of the ossification pattern in flat bones, its biochemistry is straightforward and similar to that encountered in the formation of long bones. At the nanoscale calcium phosphate is incorporated into collagen fibers to generate mineralised bone such that organisation of collagen fibers determines availability of nucleation sites [9, 20, 21]. The complex boundary condition at the growing edge of the bone and the co-occurring cellular events that drive morphogenesis (such as division, and differentiation) with asynchronous timing will, thus, generate the complexity of ossification patterns during development. Therefore, constructing a simple and random generative model of ossification should aid in understanding the formation of the ossification patterns. In our model, the intensity at lattice site (*i, j*) corresponding to the ossification level there increases by one unit with a probability *P* (*i, j*) (Fig. 1E) that incorporates three effects: a phosphate gradient, positive local feedback, and a collagen fiber field. The first effect is captured by considering a one-dimensional exponential (as shown in Fig. 1F) multiplied into the other two terms. This recapitulates the necessity of phosphate, and results in a gradient of ossification intensity. The second effect is included to capture the fact that the ossified points facilitate mineralization in their neighborhood, and set a positive feedback. This is done by convolution of the image with a box or a Gaussian kernel such as the one shown in Fig. 1G. Finally, the last effect is incorporation of minerals around the collagen fibers which is implemented here by considering an additional probability around the fibers as shown in Fig. 1H. For more details, see Supp. G. We first assumed that collagen fibers are randomly placed with uniform distribution of their centers across an area corresponding to a section of the ossification data with size 1500 *×* 900 *µ*m^2^. We furthermore assumed random orientation of fibers with uniform distribution between 0 and *π*. A typical result of this model is depicted in Fig. 1I next to a segment with the same size from WT data from E15.5 shown in Fig. 1J.

To quantitatively compare our simulation results to the experimental data, and to describe how the mineral pattern evolves over developmental time, we then sought to exploit the concept of self-similarity based on scaling behavior of given statistics of the data. As collagen comprises hierarchical repeating patterns that form the foundation of mineralized bone [8], we employed multifractal analyses for quantifying pattern maturation. Amongst many tools developed for characterizing multifractality of stochastic time-series, Multifractal Detrended Fluctuation Analysis (MFDFA) stands out due to its simplicity as well as its accuracy when applied to small data sets [18, 22]. This method can also be generalized to study two-dimensional structures e.g. common gray-scale images [23]. After segmenting the image into non-overlapping squares of a given size *s* (Fig. 2A step 1), each segment is integrated (i.e. cumulative summation in both directions is calculated as shown in Fig. 2A step 2). One then needs to detrend the data in each segment by fitting a polynomial to the cumulative sum and calculating the residuals (Fig. 2A step 3). The second-order moment of residuals in the segment is called the detrended fluctuation representing the fluctuations around the trend in each segment. We next explore the scaling behaviour of the so-called fluctuation function *F* (*q, s*) which is the *q*th-order moment of these detrended fluctuations as the size *s* is varied across the relevant range (Fig. 2A, step 4). We can then extract the scaling exponent of *F* (*q, s*) known as generalized Hurst exponent *h*(*q*), which would be a constant line independent of *q* if our signal were monofractal. Instead, we find that the Hurst exponent decreases as *q* increases, a clear indication of multifractality (Fig. 2A, step 5).

To understand what properties within our dataset give rise to this multifractality, we tested surrogates which alter one or more properties of the image while preserving the others. We generated the Ranked-Wise (RW) Gaussian, random phased Gaussian, and shuffling surrogates [24] and then analyzed them to compare with typical images from E15.5 (shown in Fig. 2E step 5) and E16.5. We found that the generalized Hurst exponent *h*(*q*) of the RW Gaussian surrogates is closest to that of our original data. It should be noted that this surrogate changes the distribution of intensities of pixels to Gaussian while preserving all linear and non-linear correlations. Further, removing non-linear correlations using the random phased surrogate results in a monofractal image with a constant *h*(*q*) equal to the offset of the original *h*(*q*). Finally, shuffling the pixel locations, which removes all correlations but preserves the distributions, results in a monofractal image which has the most different generalized Hurst exponent. Together, these results indicate that the long-range non-linear correlations are the source of multifractality in mineral pattern (see App. D for more details).

To understand the dynamics of these patterns in developmental time, we chose to extract a quantitative metric to compare self-similarity across samples from different developmental stages. Therefore, we transformed the generalized Hurst exponent into a singularity spectrum, *f* (*α*) which is simpler to interpret, using the Legendre transformation:

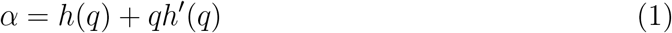

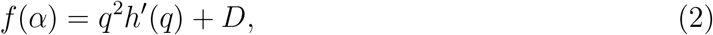

where *D* is the space dimension (*D* = 2 in this case). *f* (*α*) is the Hausdorff dimension of all data points with the given singularity exponent *α*. A rigorous and formal definition for singularity exponent *α* is given by the *Hölder* condition [25] (as discussed in App. E). Equivalently, singularities can be interpreted as topological defects with diverging magnitude in the *n*th-order gradient field of the grey-scale intensity (see App. E). As an example, mineralisation foci in developing bone are anisotropic, potentially resulting in local discontinuities in various orders of the gradient fields [9, 12, 26]. One would therefore expect to find singular points of varying strength in ossifying bone. A typical singularity spectrum is depicted in Fig. 2B from which multiple measures can be defined for characterizing the multifractal properties of the data. It should be noted that a large singularity exponent *α* corresponds to a more regular singularity with a greater number of well-defined derivatives. For example, consider a singularity with exponent *α* in a grey-scale image. Intensity at that point has *n* well-defined derivatives for *n* the largest integer smaller than *α*. Meanwhile, the Hausdorff dimension *f* (*α*) determines how space-filling a type of singularity is. *f* (*α*_max_) = 2 means that singularities with exponent *α*_max_ are found everywhere in the 2D image while other singularities form a fractal with the fractal dimension of *f* (*α*) *<* 2. As one measure of the degree of multifractality, we consider the width of the spectrum Δ*α* := max(*α*) *−* min(*α*). We also make use of the minimum *α* as a complementary measure, which represents the irregularity of the most irregular singularities in the data. We compared these two measures across developmental stages (E14.5, 15.5, 16.5, 17.5, and 18.5) and found that the variance of Δ*α* narrows over time (Fig. 2C) whereas the min(*α*) distribution has nearly constant width with an apparently oscillatory average (Fig.2D). The former behavior suggests greater subject-to-subject variability in early stages which becomes more regular and robust over the course of mineralization.Further, progressive space-filling in the mineral pattern may indicate distinct phases where early mineral has a loose pattern and mature mineral is increasingly continuous. This could be because the mineral pattern is less complex in the bone center compared to the leading edge, indicating that the bone center is more mature than the bone front (also suggested by [14]). We studied the behavior of other characteristics of the singularity spectrum as well (Supp. F). However, their dynamics are all similar to either of the typical behaviors already observed in Δ*α* or min(*α*).

Having characterized the experimental data based on the associated singularity spectrum, we applied MFDFA on our simulation results to unveil the contribution of different processes involved in ossification to the overall multifractality. We performed a sweep over all parameters in our model (Supp. G) and found that the most crucial parameter for constructing multifractal ossification patterns is collagen density. At low densities, the patterns show a high level of multifractality through Δ*α* := max(*α*) *−* min(*α*) as shown in Fig. 2E. The purple circles here show where the multifractality crosses 2 which is approximately the mean value for the images from E15.5. Moreover, the black line shows the curve that can be derived from scaling arguments connecting the fiber number to length (Supp. G 1). It should be noted that at high fiber density, no multifractality was observed independent of the other parameters, suggesting the criticality of the fiber density. We then studied the effect of having a gradient of collagen fibers by assuming an exponential probability distribution with a given length scale *γ*. Fig. 2F shows the effect of *γ* as well as *β* which is the length scale of the phosphate gradient. Longer length-scales corresponding to flatter gradients result in less multifractality, demonstrating the importance of these gradients in producing multifractal features.

To further investigate the relation between multi-fractal singularities and ossification patterns, we sought to find the spatial distribution of these singularities in the images. The Wavelet Transform Modulus Maxima Maxima Method (WTMMMM) can detect singularity points in space and determine multifractal characteristics of data. The conceptual basis of singularity detection methods is depicted in Fig. 3. Fig. 3A shows a simple example containing an isolated singularity and a local maximum. In order to find the location of the singularity, one can define a measure (*e*.*g*. sum) over a neighborhood of a given size for each point in the data, then vary the neighborhood size and observe the scaling behaviour of the measure. Over a range of small scales, regular points have arbitrary exponents determined by the properties of the measure, whereas singularities have their *own* exponents, reflecting their irregularity. As an example consider the sum measure, depicted in Fig. 3B for the simple case from Fig. 3A. The power-law behavior of three points (the singularity, maximum and a random point in the field) are shown in Fig. 3D. As one can see here, only the singularity shows non-arbitrary exponent (*i*.*e. ≠* 2). In WTMMMM –a more advance method of singularity detection–, the measure is defined as the gradient of the wavelet transform as shown in Fig. 3C. In other words, the image is smoothed by the transform and the rate of the change of the intensity of the smoothed image is used as the measure for singularity tracking. One benefit of WTMMMM is that here only maximum values of the measure are stored and analyzed to detect the singularities. Therefore, analyzing large images can be done with high precision, high speed, and with less memory. Fig. 3E, shows the performance of WTMMMM for determining the isolated singularity in Fig. 3A. Here, Gaussian wavelet is used, resulting in an arbitrary exponent of 1 at non-singular points. However, the exponent of the power-law behavior of the measure equals the exponent of the singularity. By employing WTMMMM, we detected the singularities and their exponents in typical images from E15.5 (Fig. 3F).

**FIG. 3:**
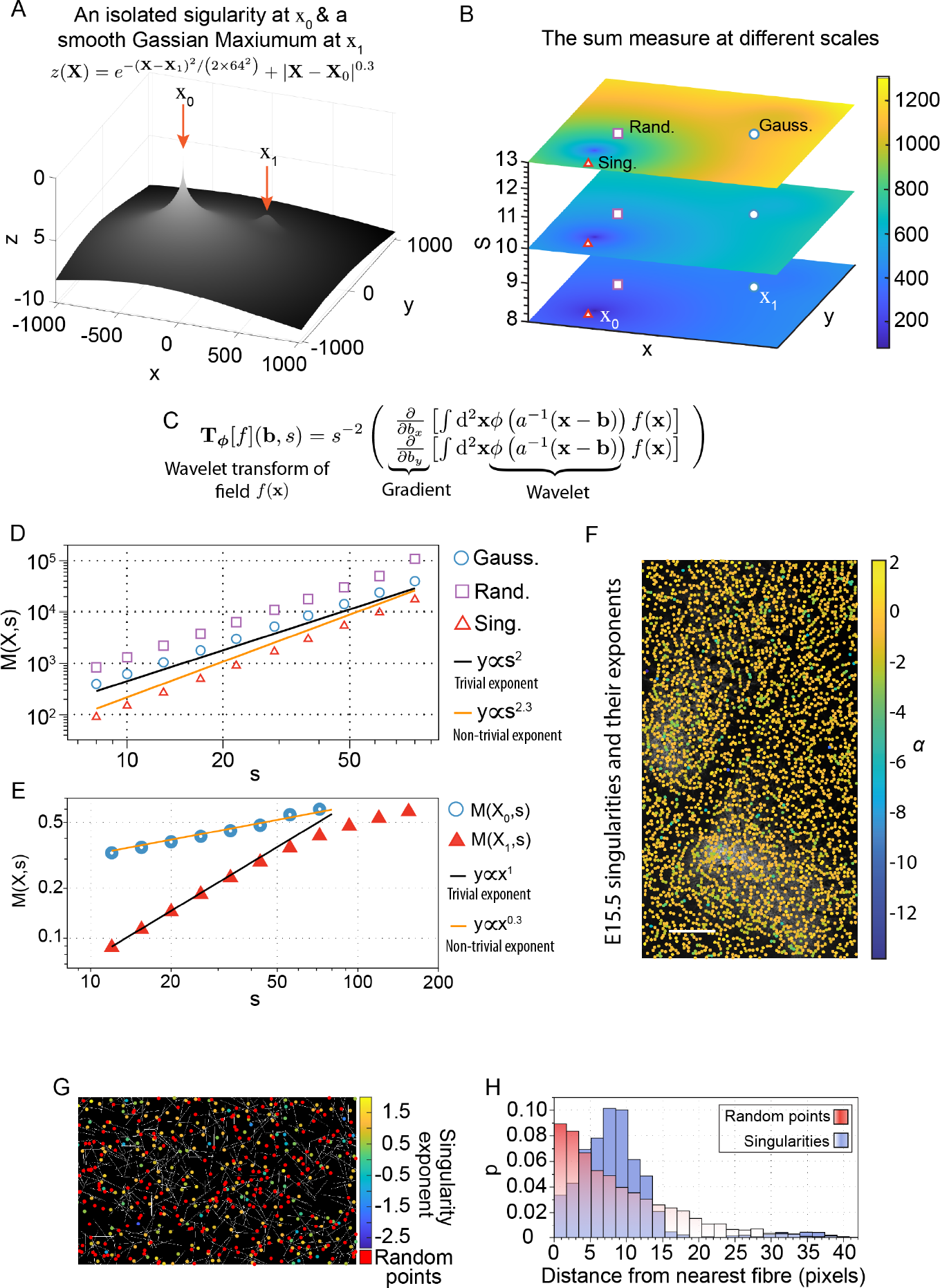
Singularity detection and singularities in ossification patterns. **(A)** A simple field containing an isolated singularity at *X*_0_ and a local Gaussian maximum at *X*_1_. **(B)** Smoothed version of that field by the sum measure at different scales. Here, the red trianles represent *X*_0_, blue circles show *X*_1_ and the purple squares are a random point in space. **(C)** The wavelet transform of the data *f* (*X*) with wavelet *φ* with scale *s* at point **b. (D)** The power-law behavior of the sum measure for three different points highlighted in (B). The exponent of the non-singular points is determined by the measure (equal to 2). **(E)** The power-law behavior of the wavelet transform modulus with Gaussian (first-order) wavelet for the maxima and singularity of A. The exponent of the Gaussian maximum points is determined by the wavelet order which is 1 here and for the singularity is 0.3. **(F)** Singularity exponents and their locations detected by WTMMM for WT E15.5. We here use a Mexican hat wavelet which is capable of detecting singularities with exponents up to 3.**(G)** A typical collagen fiber field (white lines) used for ossification simulation along with the singularities detected via WTMMMM with the Mexican hat wavelet (shown by color coded circles) and random points (shown by red circles). **(H)** The histogram of distances between singularities and their nearest fibre (in blue) and that of random points (in red). The singularities are mostly located at a certain distance from fibres, corresponding to the wavelets used for the analysis, while distances of random points have an exponential distribution, a signature of Poisson processes.

So far we have focused on the global multifractal features produced by our generative model. However, useful insight can be gained by studying the location of the singularity points in the patterns as well. We applied WTMMMM on the simulation results and detected singularities with exponents ranging from -2.7 to 1.8. Fig. 3G shows a typical distribution of singularities (color coded circles) compared to collagen fibers (white lines). As one can see, the singularities co-localize with the fibers. To better represent this we also distributed the same number of points randomly with uniform distribution (shown by red circles), and found the distance of every point to the nearest fiber. As shown in Fig. 3H, the distances between singularities and fibers have a peak around 8 pixels corresponding to the smallest scale used in WTMMMM which can be interpreted as the effective resolution of the method. However, the distribution of distances from random points to the fibers follows an exponential. This is consistent with production of singularities as a result of nucleation centers.

Our toy model predicts that phosphate and collagen gradients together with a collagen density *below* a critical value are necessary to recapitulate multifractal patterns found *in vivo*. To test these predictions we sought to perturb collagen organization in such a way that would restrict phosphate incorporation and alter collagen distribution across biological scales within the tissue. Emergent collagen structure in tissues such as bone requires crosslinkers that cannalise the physical association of individual fibers through which phosphate can be incorporated. To perturb this collagen structure and its distribution we chose to inhibit Lysyl oxidase (LOX), a predominant Col1a1 crosslinker, using an irreversible LOX inhibitor, Beta Aminoproprionitrile (BAPN) [27]. After feeding pregnant dams 0.1% BAPN treated or control food from the beginning of skull development (E11.5), we collected skull caps at E15.5 and E17.5 marking the middle and end stages of bone morphogenesis. Indeed, we found qualitative differences in mineral pattern of BAPN-treated skulls where the mineral pattern exhibited greater lace-like characteristics at E15.5 as one can see in Fig. 4A,B. *In vitro* mineralization experiments suggest a spatial relationship between collagen fibrils and phosphate incorporation mineral nucleation that occurs in close proximity to collagen fibrils [9, 12, 26]. We also observed colocalization of singularities in ossification patterns and collagen fibers in our *in silico* experiments. If so, we would find an altered pattern of mineral foci within BAPN-treated skull caps. Transmission Electron Microscope tomograms of E15.5 skull caps revealed an increase in the size of mineral foci and variability in foci shape within BAPN-treated bone as shown in Fig. 4C-D. These data confirm that nano-scale collagen structure and organization translate into mesoscale mineral pattern. Indeed, upon applying MFDFA to our BAPN samples at the end of morphogenesis we found a significant increase in the min(*α*) of the patterns indicating an increase in the regularity of the singularities. To further explore these data we investigated the spatial aspect of multifractality in control and BAPN-fed animals and mapped singularity points to our images using our WTMMMM at E17.5 (see Fig. 4H, I). As one can see, the more irregular singularities (i.e. those with more negative exponents) in the WT image cluster together. This is better captured by Fig. 4J which depicts the distributions of distances between singularities with exponents *α ≤ −*3. Indeed, the distances amongst the irregular singularities in WT patterns have an extra peak at shorter distances, corresponding to clustering. Recall that the phase diagram rendered from our generative model predicts the presence of a strong decrease in Δ*α* for increasing fiber densities, as the sharp transition line in the parameter space is crossed. As the Δ*α* of BAPN treated samples are consistent with such an increased fiber density, we asked whether BAPN induced an increase in collagen production that would generate increased fiber density within the bone *in vivo*. qPCR indeed revealed a significant increase in the relative expression of Col1a1 in BAPN-treated samples compared to Control fed animals as presented in Fig. 4E. By increasing the density of collagen by a factor of 2, the model predicts a near total collapse of multi-fractality, as measured by Δ*α*, and indeed the measured Δ*α* in the BAPN treated samples quantitatively matches this prediction Fig. 4K.

**FIG. 4:**
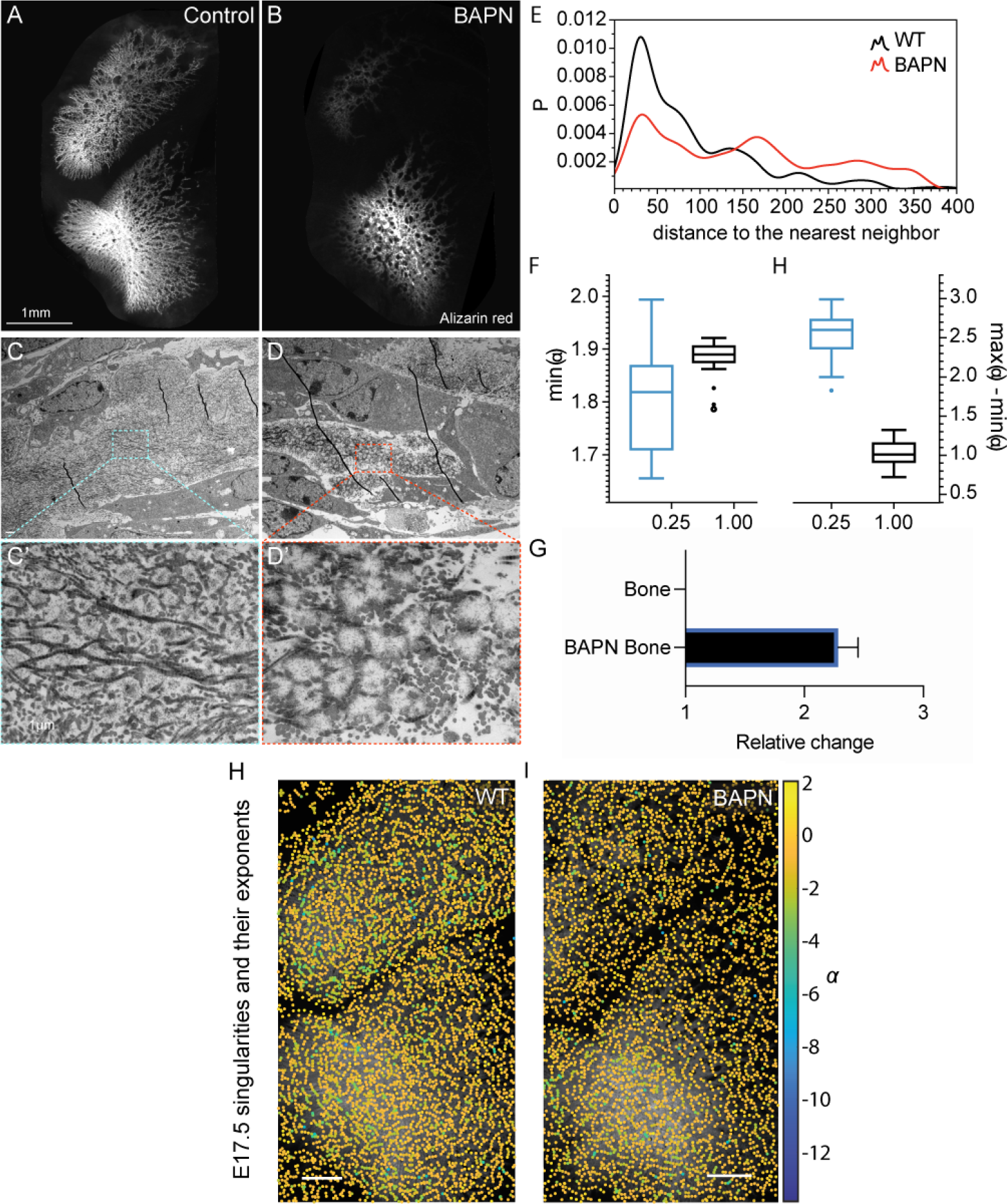
Perturbation of collagen fiber organisation. **(A)** control ossification pattern from E15.5. **(B)** BAPN-treated ossification pattern from E15.5. **(C)** Transmission Electron Microscope tomograms of WT E15.5. **(D)** Transmission Electron Microscope tomograms of BAPN-treated subjects from E15.5. **(E)** Comparison of multifractality Δ*α* in WT and BAPN-treated subjects from E15.5. **(F)** Comparison of irregularity of the most irregular singularities min(*α*) in WT and BAPN-treated subjects from E15.5. **(G)** Relative changes in the abundance of collagen1 in WT and BAPN-treated subjects from W17.5. **(H)** and **(I)** Singularity exponents and their locations detected by WTMMM for WT and BAPN from E 17.5, respectively. We here use a Mexican hat wavelet which is capable of detecting singularities with exponents up to 3.

### Discussion

We believe that the power of this study lies in the successful marriage of a simple model with few assumptions to sophisticated analysis of complex data. For example, our model predicts that higher fiber numbers have a greater capacity to suppress multifractality than can be compensated for by shorter fiber lengths. This was verified experimentally by our BAPN-mediated perturbation of collagen structure as BAPN changes both the size and aspect ratio of fibers (see [28]. A further prediction that arises from combining our model, data, and analysis in this way, is the existence of a spatial gradient of collagen across the tissue. Indeed, recent work from the Tabler lab demonstrates the generation of a collagen gradient during these stages of skull expansion [14].

These predictions and insights into the tissue are critically made possible by the advancement of multifractal analysis techniques presented here. Traditionally, monofractal analyses due to their limitations have been used to describe restricted areas such as spatial networks or bone but rarely consider the possibility that the systems under study are multi versus monofractal. This can occur because only one statistic is often considered and a range of scalings cannot be detected from evaluating only one statistic. This work demonstrates firstly, that monofractal assays are insufficient to describe ossification patterns in tissues, despite their popularity in diagnostic techniques. Secondly, this work, therefore, opens the door to a fuller understanding and greater analytic power across a wide swathe of other mesoscale biological patterns. Many tissues that may appear not to be fractal in nature could harbor multifractal features that can be used to describe pattern and add to biological insight of how these patterns form. In many cases, mesoscale patterns are believed to emerge from interactions at multiple lower scales. Examples of multiscale interactions implicated in pattern formation are myriad and include tissues such as the Purkinje fibers of the heart [29], the hepatic system [30], or insect chitin [31]. In such cases, generically one finds different properties scaling in distinct ways, which is precisely the optimum usage of multifractal analyses.

At its core, multifractal analyses like those presented here provide access to spatial variations that are typically lost by averaging. Even in the absence of averaging, the focus on one particular feature within the data prevents a holistic understanding of the process and potential links between features. Indeed here, we first sought to understand the spatial distribution of voids and the ossified bone they reside within. By implementing these analyses, we developed more than a quantitative metric of mineral pattern and could generate new hypotheses which we later tested *in vivo*.

Here we show a work program through which one can leverage the complexity common to morphogenetic and late-stage developmental data. Where live imaging is often not possible our approach provides an opportunity to infer dynamic features of emergent pattern from static images. Meeting that complexity head-on with our multifractal approach which does not average out statistics of spatial pattern, we have shown how one can implement a complete loop of experiment, analysis, and modelling without the use of *a priori* explicit mechanistic knowledge. Yet, our approach has yielded a multiscale mechanistic insight that would otherwise have been inaccessible. However, much remains to be understood not just in bone but broadly in the emergence of developmental systems.

## ACKNOWLEDGMENTS

We are happy to acknowledge S.W. Grill, R. Mateus, and sT.H. Tan for a close reading and helpful feedback on the manuscript. We are also particularly grateful to S.W. Grill for a further number of invaluable discussions. We would like to thank the following Services and Facilities of the MPI-CBG for their support: Electron Microscopy, Light Microscopy, and Biomedical Services. M.B. was partially supported by the German Federal Ministry of Education and Research under grant number 031L0160. M.B. was subsequently further supported by the European Union’s Horizon 2020 Research and Innovation Programme under grant agreement no. 829010 (PRIME) during the completion of the manuscript.

## Appendix A Materials and methods

### Mouse lines

C57Bl/6JOlaHsd or C57Bl/6NTac mice were used to generate skull cap images and EM. BAPN Experiments conducted at the MPI-CBG adhered to the German Welfare Act and were overseen by the Institutional Animal Welfare Officer.

### Lysyl-Oxidase Inhibition

For collagen crosslinking inhibition studies, pregnant females were fed 0.25% (1 g*/*kg) BAPN containing food from E11.5, the onset of skull morphogenesis. For embryo collection, pregnant females were euthanized by cervical dislocation, and embryos were collected for downstream analysis. Protocol was approved by the Institutional Animal Welfare Officer. BAPN containing food was made by Safe Complete Care Competence.

#### 1 Alizarin red staining

After fixing Embryos in 4%PFA overnight at room temperature embryos were placed in 1xPBS and extraembryonic membranes were removed. Embryos were then fixed in 95% ethanol for 1 hour. Embryos were then placed in acetone overnight at room temperature. Acetone was then replaced with Alizarin red staining solution (0.005% in 1%KOH - 10mg Alizarin red in 200ml 11% KOH) and embryos were incubated on a rotating shaker for 3 hours. After, embryos were washed overnight in 1% KOH and then transferred to 50% glycerol: 50% 1%KOH solution and incubated at room temperature until tissue appeared transparent. Once cleared embryos were transferred to 50% glycerol: 50% ethanol.

### Flat-mount skull cap imaging of stained samples

After ataining, skull caps were mounted flat in 50% glycerol: 50% ethanol and imaged using the Zeiss AxioZoom ApoTome system.

#### 2 Transmitted Electron Microscopy

WT and BAPN embryos were collected as described above and fixed with 5% glutarladehyde/ 1% tannic acid in 0.1M PBS pH 7.2. Fixed heads were embedded in 4% low melting agarose and 200 µm sections were obtained using a vibratome (Leica, VT1200S). Sections were post-fixed with 1% osmium tetroxide (Electron Microscopy Sciences; Cat# 19190) in water. Sections were dehydrated in serial steps (30%, 50%, 70%, 80%, 90% and 100%) of Ethanol (EtOH), infiltrated with 1:3 EPON LX112/EtOH, 1:1 EPON LX112/EtOH, 3:1 LX112/EtOH and pure LX112. Sections were embedded on Teflon-coated slides with Aclar spacers (7.8 mil, Science Services and Miller-Stephenson; ordering number: DF-3R, respectively). Ultrathin cross sections (70 nm) were obtained using an ultramicrotome (Leica, FC7). Sections were post-stained with uranyl acetate (Electron Microscopy Sciences; Cat# 22400) and lead citrate (Electron Microscopy Sciences; Cat# 17800) and viewed in a Morgagni (FEI/Thermoscientific) transmission electron microscope, equipped with a Morada CCD camera (Emsis), at 80 kV.

## Appendix B Multi-Fractal Detrended Fluctuation Analysis in two dimension

The basic idea of MFDFA in two dimension is discussed in the main text, but the detailed description of steps are presented here:

- **Step 1**. Divide the surface under study (i.e. the intensity of the image) given by a two-dimensional array *X*(*i, j*) with size *N × M* into *N*_*s*_ *× M*_*s*_ disjoint boxes of size *s × s* (see Fig. 1E). Here, 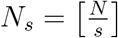 and 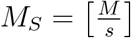. Each segment can be represented by *X*_*v,w*_(*i, j*) = *X* ((*v −* 1) *s* + *i*, (*w −* 1) *s* + *j*) where *i, j* = 1, 2, …, *s*.
- **Step 2**. Calculate the cumulative sum *Y*_*v,w*_ for each segment *X*_*v,w*_ via the following relation:

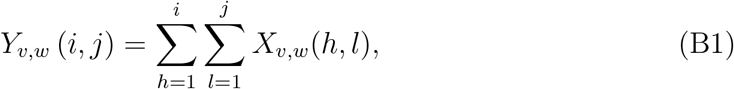

where *i, j* = 1, 2, …, *s*. In fig. 1E step 2, one can see the cumulative sum shown by red points.
- **Step 3**. Detrend the cumulative sum surface in each segment *Y*_*v,w*_ by first fitting a bivariate polynomial function 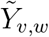 and then obtaining the residuals

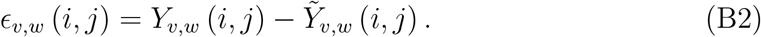

Then, calculate the “detrended fluctuation” at segment *v, w* as

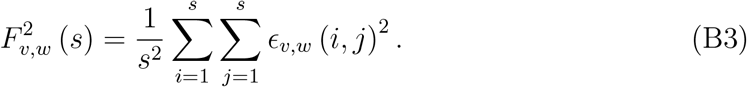

In fig. 1E step 2, the polynomial fit to the cumulative sum is shown by the solid plane, and in step 3, the residuals are shown. The procedure of choosing the order of polynomial fitted in eq. B2 will be discussed later in the remarks section.
- **Step 4**. Calculate the q-th order overall *fluctuation function*

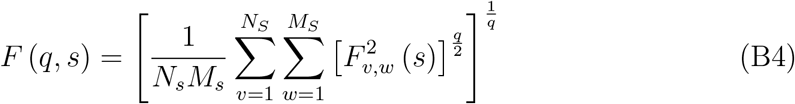
- **Step 5**. Determine the scaling behavior of the fluctuation function *F* (*q, S*) by varying *s* for each value of *q* reading

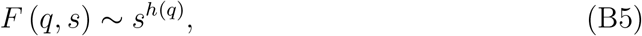

where *h* (*q*) is generalized Hurst exponent. This can be obtained by simply fitting a line to the *log − log* plot of *F* (*q, s*) vs. *s* for each value of *q* i.e.

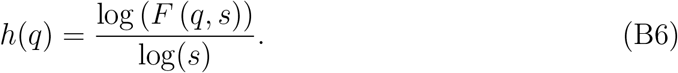

In step 4 of fig. 1E, one can see the scaling of the fluctuation function *F* (*q, s*) for a few *q* values. The generalized Hurst exponent for a typical ossification pattern is shown in fig. 1E step 5.

It should be noted that the traditional DFA method can be realized by performing steps 1–3 exactly as mentioned above and the remaining steps with *q* = 2. In this way, only the Hurst exponent *H* = *h*(*q* = 2) can be determined.

### 1 Choosing the parameter for MF-DFA

The parameters that should be used for MF-DFA vary from one system to another depending on their properties and therefore they should be determined separately for each data set. In this study, we used the following procedure for a small subset of samples (from stage 15.5D) to determine the parameters and then used these parameters to analyze all images.

In order to determine the range of scales *s* over which the analysis is done, one needs to start from the widest meaningful range. This range should start from the order of the fitted polynomial, so that the number of parameters does not exceed the number of points. Moreover, the range of *s* should stop at *s*_max_ that results in a number of segments being large enough so that the averaging process in eq. B4 remains meaningful. Once this maximum range is determined, one should find the sub-range in which the fluctuation function *F* (*q, s*) shows power law scaling for all values of *q*. This range can be then used for the rest of the analysis.

In eq. B4, *q* is the order of the moment of the fluctuations. The range of this parameter should be chosen according to the minimum number of segments. There should be high number of segments for each value of *q*. Large values of *q* correspond to statistics of the segments with large fluctuations and negative values correspond to the segments with small fluctuations. Therefore, maximum and minimum *q*’s should be chosen in a way that their corresponding fluctuation functions behave similar to others, i.e. show power law behavior over the chosen range of *s*.

The order of the chosen polynomial trend depends on the nature of the data and any intrinsic trends contained therein. As described in [19], we determined this order by starting from the simplest polynomial 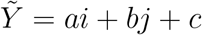 and increasing the order in *i* and *j* until the surface *Y*_*v,w*_ (*i, j*) is over-fitted and the generalized Hurst exponent is no longer a monotonic function of *q*. Here, it is specially loosing its accuracy in small *q*’s corresponding to small fluctuation segments.

## Appendix C Periodic trends and frequency analysis of the data

Although MF-DFA is capable of determining multifractal features in the presence of many forms of trends in data, it fails when applied to the data sets with periodic trends. However, it is shown that such trends only cause an increase in the overall fluctuation function at scales near the wavelength of the trend which in turn results in inaccurate determination of generalized Hurst exponent [32]. One can circumvent this problem by filtering out the low frequency (i.e. long wavelength) components of the Fourier transform. This can be done by simply Fourier transforming the data, replacing the low frequency terms by zero one by one, then taking the inverse Fourier transform, and analyzing the result until the crossover is removed.

In Fig. 5a, the overall fluctuation function *F* (*q, s*) is shown for a typical ossification at stage 15.5 with *q* ∈ {−3, 2, 1, 0, 1, 2, 3}. As one can see in this plot, there is a sudden increase in *F* (*q, s*)’s at the large values of *s*. This crossover is more pronounced for smaller *q*’s but exists in all of them which makes determination of *h*(*q*) inaccurate. By removing a few of the low frequency components of the data, however, one gets proper power laws within the same range of *s*. In Fig. 5b, the same plot as in 5a is shown but after removing two lowest frequency components in all directions. It is worth noting that removing more frequency components results in flattening the generalized Hurst exponent i.e. suppressing the multi-fractality. In Fig. 6, one can see the generalized Hurst exponent *h* (*q*) of the data whose results are shown in Fig. 5 with various number of low frequencies (*m* ∈ {0, 2, 5, 8, 11, 14, 17}) removed. As one can see here, the overall multi-fractality which can be defined as the change in *h*(*q*) decreases as we increase *m*. Therefore, we only use *m* = 2 for analyzing all images in order to prevent over-filtering and getting the most accurate multi-fractal measures. A similar behavior has been observed in another context where low-pass filtering (removing the highest frequencies) also results in flattening of *h*(*q*) when analyzing time-series recorded from V1 cortex [19].

**FIG. 5:**
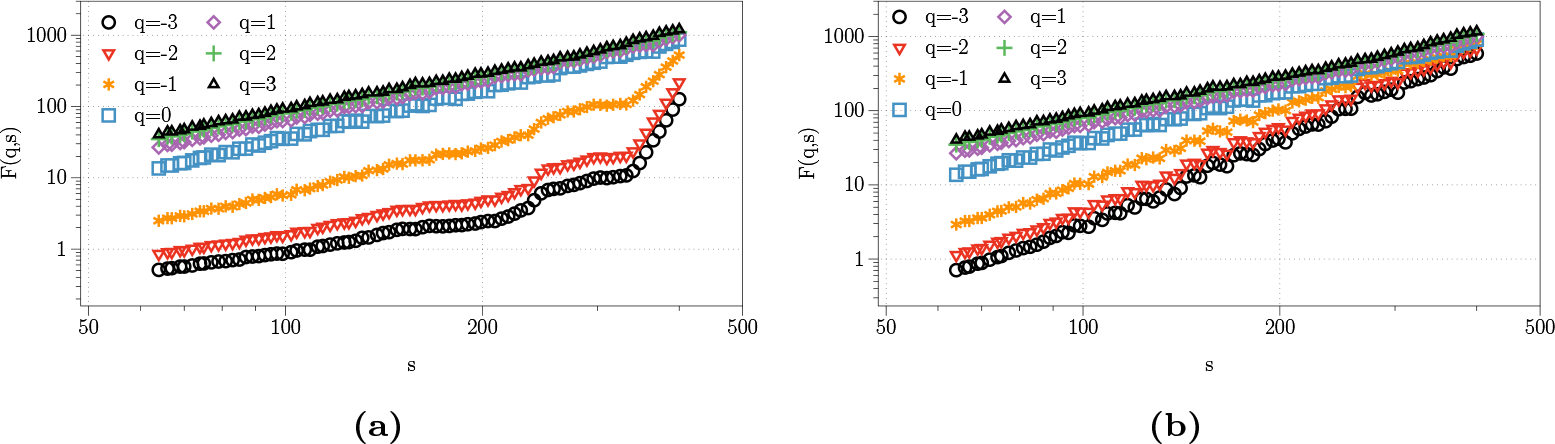
**(a)** *F* (*q, s*) vs. *s* for a typical and raw image at 15.5 stage. **(b)** *F* (*q, s*) vs. *s* of the same image after removing 2 of the lowest frequencies in all directions.

**FIG. 6:**
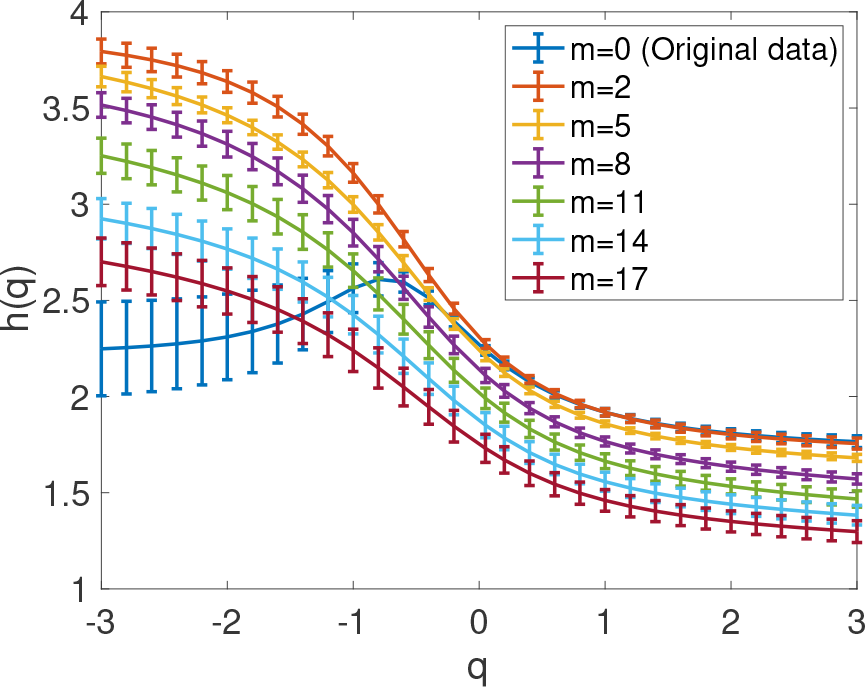
Generalized Hurst exponent of a typical ossification pattern after removing *m* lowest frequency.

## Appendix D The origin of multifractality in the data

A data set can exhibit multifractality for various reasons including: I) non-Gaussian (e.g. fat-tailed) distributions, II) existence of linear or III) nonlinear long-range correlations. One can investigate the contribution of each of these properties to the overall multifractality using surrogates which change one or more properties while preserving the others. Here, we first use Ranked-Wise (RW) Gaussian surrogate [24], which preserves all linear and non-linear correlations but changes the distribution of the intensities to Gaussian. We also use random phased surrogate to preserve linear correlations, remove nonlinear correlations, and change the distribution to Gaussian with the help of the central limit theorem. Finally, we shuffle the pixels in the image to preserve the distributions while removing all correlations.

We analyzed two images from 15.5 and 16.5 stages, and as one can see in Fig. 7, the closest curve to the *h*(*q*) of the original data is the RW-Gaussian surrogate. The one which differs the most is unsurprisingly the shuffled one, which only preserves the distribution. One can accordingly conclude that the non-Gaussian distribution has the least contribution, and the long-range correlations contribute most to the overall multifractality. Moreover, one can consider a random-phased surrogate and argue that the linear correlations set the offset of the generalized Hurst exponent *h*(*q*) but do not contribute significantly to the multi-fractality. We therefore established that the observed multifractality originates primarily from *long-range non-linear* correlations in the data.

**FIG. 7:**
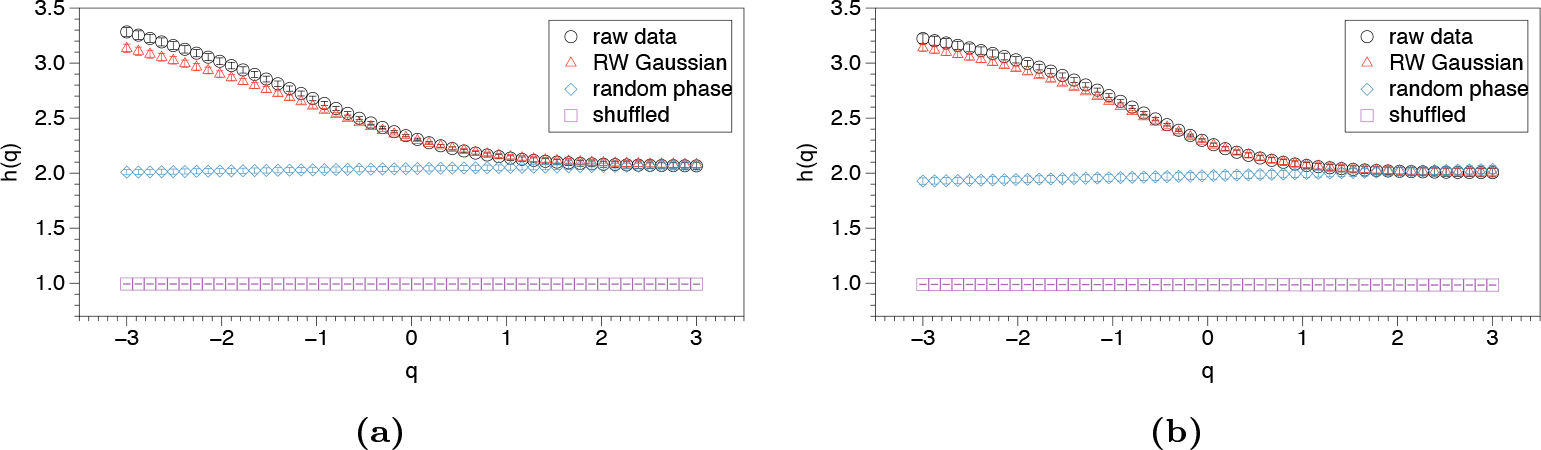
Investigating the origin of multifractality in our data from **(a)** E15.5 and **(b)**E16.5

## Appendix E Singularities and singularity exponents

As discussed earlier, the direct output of MF-DFA is the generalized Hurst exponent *h*(*q*). This quantity represents the scaling behavior of *q*th-order moment of fluctuations in the data. The generalized Hurst exponent *h*(*q*) can be transformed into the singularity spectrum *f* (*α*) via the Legendre transform

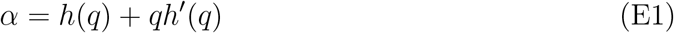

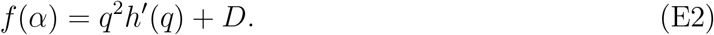

This spectrum is generally easier to interpret and more useful to characterize the data. Here, singularity exponent *α* (also known as H older exponent) determines the regularity of the singularity. To be more precise, consider function *f* (**r**) has a singularity at **r**_0_ with exponent *α*. Then, there exist two constants *C* and *h*_0_ *>* 0, and a polynomial of order *n, P*_*n*_(**r**) with *n ≤ α ≤ n* + 1, such that

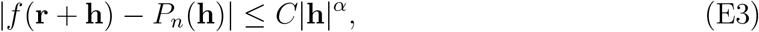

for any |**h**| *< h*_0_ [25]. This is known as Hölder condition. Furthermore, the singularity spectrum defined in Eq. E2 is the Hausdorff fractal dimension determining the fractal dimension of the set of singularity points with exponent *α*.

Although Eq. E3 provides a rigorous definition of singularity exponent, a better interpretation can be achieved by visualizing a few examples of singularities and their connection with topological defects in the gradient fields. First, note that at the singularity point **r**_0_ with exponent *α* the field *f* (**r**) is *n* times differentiable with *n* being the largest integer smaller than *α* (see Ref. [25] for more detail). Therefore, a singularity with exponents *α <* 1 results in discontinuity in the gradient field. An isolated singularity with exponent *α* = 0.5 shown is in Fig. 8A. Since for this singularity *α <* 1, one would expect a discontinuity in the first derivative which is the gradient. In Fig. 8B shows this gradient field in which an aster topological defect exist at the location of the singularity. Moreover, the magnitude of the gradient field is not well-defined and diverging at the singularity point. For a singularity with exponent *α >* 1, the largest integer *n* smaller than *α* is 1. Therefore, the first derivative is well defined and continuous everywhere. Fig. 8C shows an isolated singularity with *α* = 1.5. The gradient of this field is represented in Fig. 8D where the magnitude of the gradient field approaches zero at the singularity. However, the second-order gradient which is a tensor with four elements has discontinuity. Figs. 8E and 8F show the orientation of the eigenvectors of this tensor. Fig. 8E shows the eigenvectors corresponding to the smaller eigenvalue while Fig. 8F shows those corresponding to the larger eigenvalue. Also the absolute value of the eigenvalues are represented by the length of the lines. Here, one can see aster and vortex topological defects with magnitudes diverging again at the singularity. It is also worth noting that at local extremum points, the gradient fields have topological defects, but the magnitude of these fields either vanish or converge to a finite value.

**FIG. 8:**
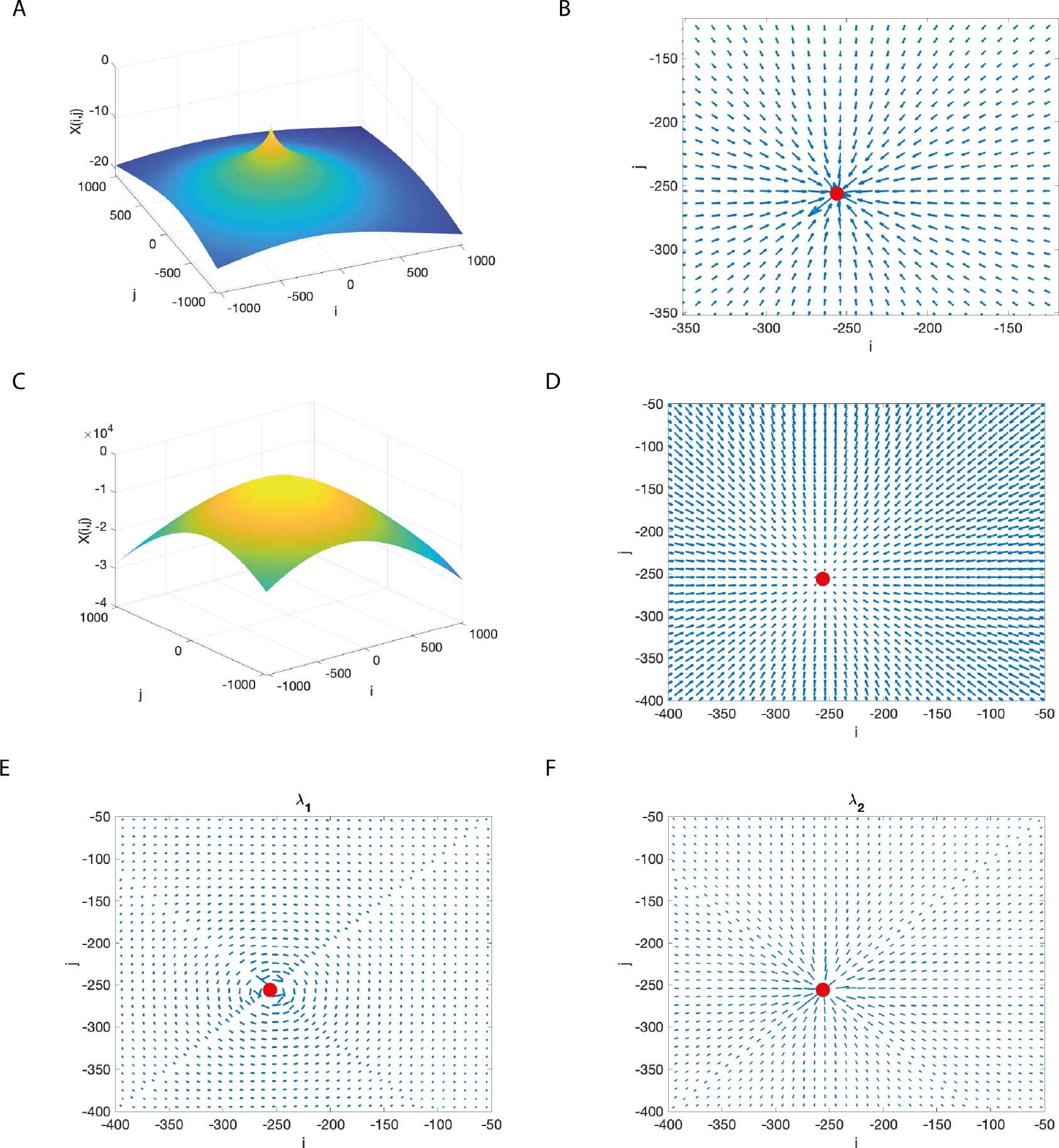
The connection between singularities and topological defects in gradient fields. **A)** An isolated singularity with exponent *α* = 0.5. **(B)** The gradient of the field represented in A. **(C)** An isolated singularity with exponent *α* = 0.5. **(D)** The gradient of the field represented in C. **(E)** The orientation of eigenvectors corresponding to smaller eigenvalue of gradient of the field represented in D. **(F)** The orientation of eigenvectors corresponding to larger eigenvalue of gradient of the field represented in D.

One can analytically study the simple isolated singularities as following: consider function *f* (**r**) = *r*^*α*^ where *r* is the norm of position vector **r**. This function has a singularity with exponent *α* at the origin **r** = 0. Let’s consider the gradient of this field

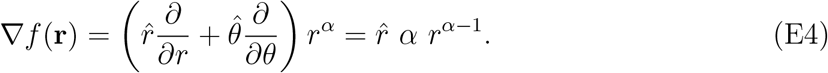

It is obvious that first of all this gradient field forms an aster at the origin. Also, the magnitude of the field diverges for *α <* 1. Therefore, there exist a discontinuity in the first gradient field if *α <* 1. However, for 1 *< α <* 2, the first gradient field is continuous as the magnitude approaches zero at the origin. In this case, the gradient of gradient can be calculated as

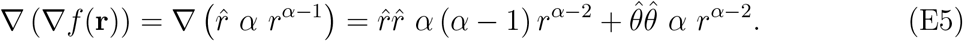

Similarly, if *α <* 2, two topological defects with diverging magnitude exist in both eigenspaces of the second-order gradient.

## Appendix F Multifractal characterization of ossification patterns over time

As mentioned in the main text, we use singularity spectrum *f* (*α*) to characterize the ossification patterns of mouse embryos during development (E14.5–E18.5). There are several properties of a singularity spectrum that one can use to characterize multifractal patterns. In this study, we used the following measures: *I)* the location of the peak *α*_max_ which is the singularity type that can be found everywhere in the data because the maximum Hausdorf dimension max (*f* (*α*)) is always 2, *II)* the minimum exponent of the spectrum min(*α*) which determines the irregularity of the most irregular singularities (see App. B for more details on the interpretations), *III)* the left part of the range corresponding to the diversity of the singularities, *IV)* the area under this part the curve which determines the abundance of the more irregular singularities, *V)* the whole range of spectra Δ*α* := max(*α*) *−* min(*α*) that represents the diversity of all singularities, *VI)* the maximum exponent of the spectrum max(*α*) which determines the irregularity of the most regular singularities and *VII)* the area under the curve over the whole range which determines the overall abundance of singularities. The bottom right pannel in Fig. 9 shows the definitions of these measures on a typical singularity spectrum. We analyzed all images from different developmental stages between E14.5 and 18.5, and Fig. 9 shows the multi fractal measures for these images. The measures I)–VII) are respectivly depicted in 9A–G. As one can see in this figure, there are two main behaviors that the measures show over time: The first behavior observed in measures I)– IV) is starting from a wide distribution and getting narrower. However, measures V)–VII) have an almost constant width of distribution, but their average change in a oscillatory way.

**FIG. 9:**
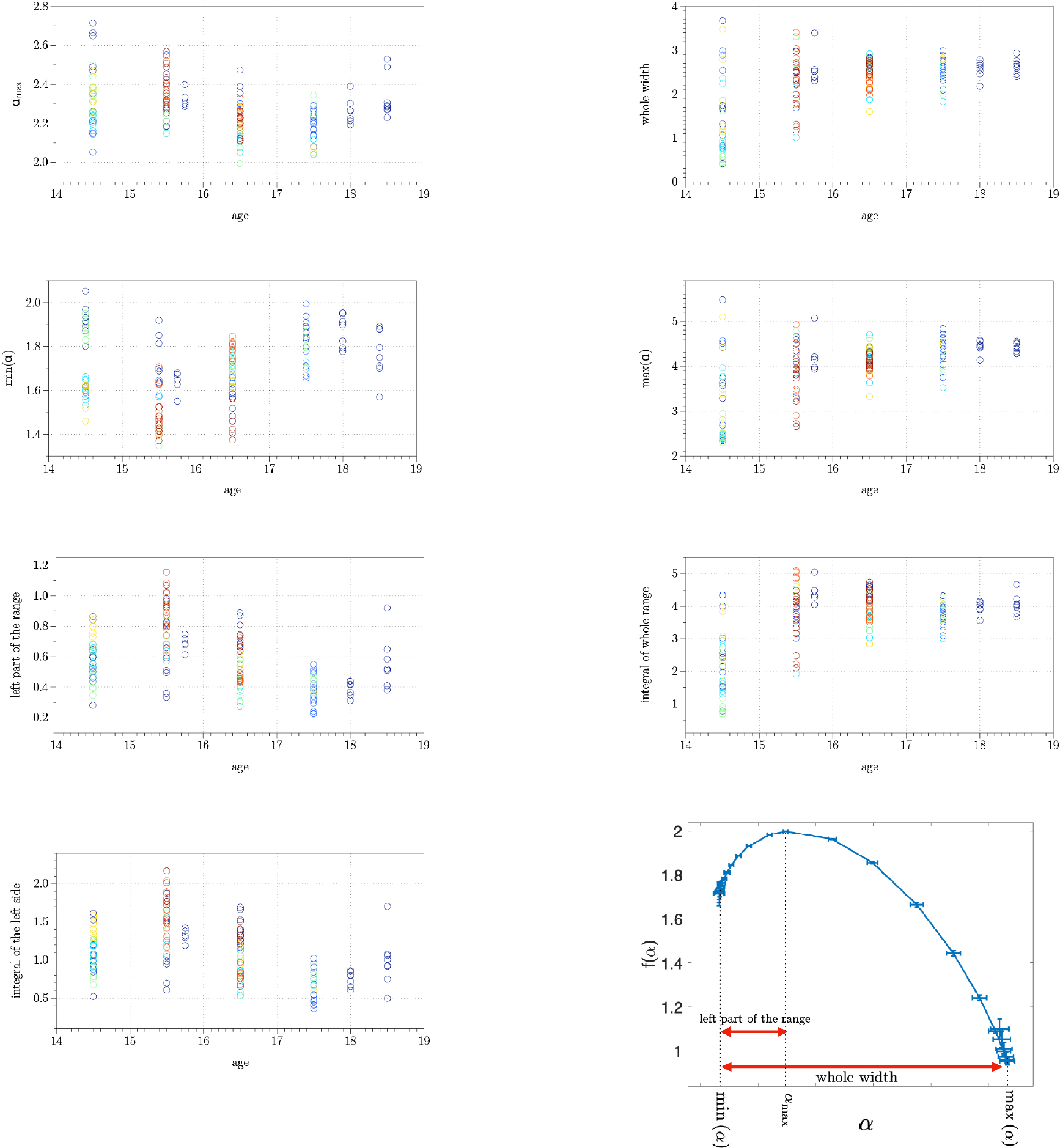
Time evolution of the multi-fractal measures. The bottom right panel shows the definition of these measures. As one can see there are two typical behaviors in these measures: left column showing an oscillation and the right column that shows a convergence to a narrower distribution.

## Appendix G Generative model

In order to further study the implications of the multi-fractal analyses, we propose a simple generative model for simulating ossification patterns which only includes the essential components of the process. We then apply the multi-fractal analyses on the results of these simulations and unveil the relation of singularities and their spectrum with the underlying mechanisms of ossification. It is worth noting that our objective here is not to reproduce the entire ossification patterns with their complicated boundary since it is a result interactions with the surrounding tissues and vasculature. We solely focus on recapitulating a small section of the ossified tissue such as the magnified pieces in Fig. 1C and D which still show multifractality when analyzed separately. Therefore, in our model, the total field is composed of 900 *×* 1500 pixels each of which correspond to a region with area of approximately 1*µm*^2^. In each iteration, the intensity of a given pixel at (*i, j*) can increase by one with the probability which incorporates three basic mechanisms known to be involved in mineral deposition [8, 33]: Phosphate metabolism which acts as an effective morphogen gradient, a positive feedback from neighboring pixels, and a field of collagen fibres that act as ossification nucleators. To be more specific, *P*_*grad*_ corresponds to the effect of the phosphate metabolism,

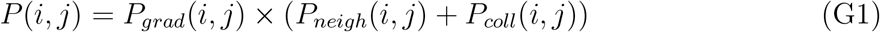

which is assumed to effectively act as a morphogen gradient. This term is proportional to the concentration of phosphate in the form of a one-dimensional exponential gradient exp (*− i/a*). It is multiplied into the other terms so that there is no ossification where this function vanishes. *P*_*neigh*_ takes into account the fact that once a pixel’s intensity is increased, its neighboring pixels are more likely to ossify and increase their intensity in the next iterations. In order to implement this, we use the convolution of the ossification intensity *X*(*i, j*) with a kernel with a given width *β* (i.e. ∑*v* ∑*u K*(*i − u, j − v, β*)*X*(*i, j*)). Here, we use box kernel for faster computations. Finally, *P*_*coll*_ is the contribution of collagen fibers such that presence of a fiber in a neighborhood increases probability of ossification. For this term, we again use convolution of a collagen field with a box kernel with width *γ*. Substituting these into Eq. G1, one gets the explicit form of the probability as

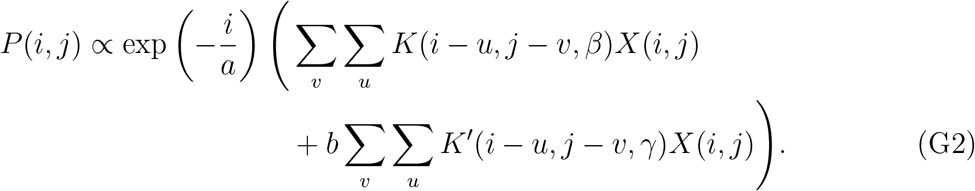

Fig. 10a shows a typical fiber field which contains 1400 fibers with length of 20 pixels randomly placed with uniform distribution. Running our simulation on this fiber template results in an image similar the one depicted in Fig. 10b if the following biologically relevant parameters are used: *a* = 30000, *β* = 20, *γ* = 12, and *b* = 7.5. Comparing this to a typical ossification pattern from E15.5 shown in Fig. 10c, one can argue that longer fibers are needed. In Fig. 10d, on can see a simulation result with very long (100 pixel long) fibers. Although, this image is more similar to the experimental data it does not exhibit any multifractality (i.e. the generalized Hurst exponent is constant). In principle, it might be possible to recapitulate both multifractality and appearance of the data by incorporating long-range interactions of fibers resulting in long persistence length. However, this type of interactions and alignment of fibers adds extra complexity which defies the purpose of our toy model. We therefore ignore the visual differences and focus on the multifractal features of our toy model.

**FIG. 10:**
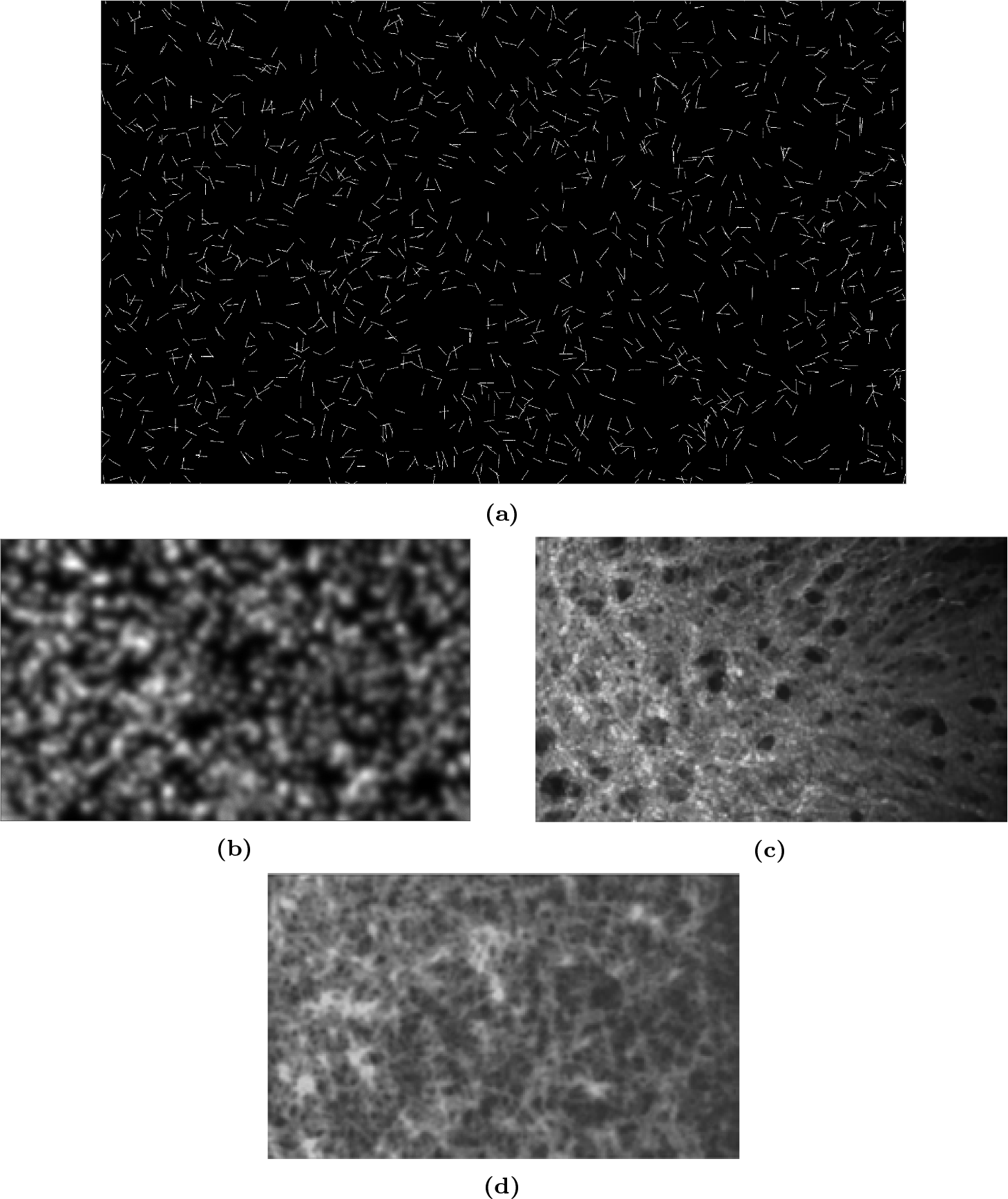
Typical example of our ossification simulations. **(A)** the collagen fiber template used containing 1400 fibers with length 20 distributed with uniform distribution across the field. **(B)** The result of our generative model for the fiber template shown in (A). **(C)** A small segment of ossification pattern from E15.5 with the same dimensions as (B). **(D)** Simulation results similar to (B) but with 100 pixels long fibers.

### 1 Determining the transition curve of multifractality using re-scaling arguments

In order to investigate the dependence of multifrcatality, measured by max(*α*) *−* min(*α*), on the density of collagen fibers, we varied the fiber number as well as their lengths. We then analyzed the results using MFDFA and the results are shown in Fig. 3G. In that plot one can define curves of constant multifractality (max(*α*) *−* min(*α*)) that suggest an inverse relation between the collagen fiber length and number. In this Sec., we use a re-scaling argument to derive an equation describing this relation.

Suppose that a box with size *L*_1_ contains *n*_1_ collagen fibers with length *l*_1_. Let us focus on a smaller box with size *L*_2_ = *λL*_1_ where *λ <* 1. Inside the new box, there are *n*_2_ = *λ*^2^*n*_1_ fibers with length 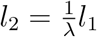. Then it is to show that

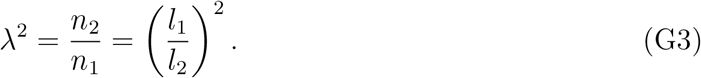

We can assume that 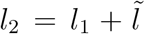 and *n*_2_ = *n*_1_ + *ñ* where the variables with tile are the differences. By substituting these into Eq. G3, one obtains

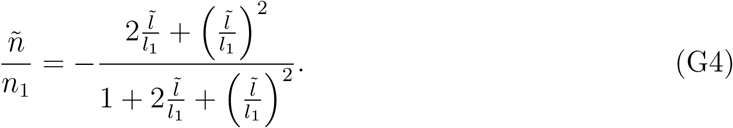

The solid black line in Fig. 3G, shows this function fitted to the points that are estimated to fall on a constant multifractality curve with max (*α*) *−* min (*α*) = 2.

## Appendix H Characterizing multifractal features using WTMMMM

### 1 The Wavelet Transform Modulus Maxima Maxima Method

The Wavelet Transform Modulus Maxima Maxima Method (WTMMMM) is based on tracking the gradient maxima of the wavelet transform moduli over a range of small scales. A detailed discussion of this and the necessary steps is provided in Ref. [34]. We here only discuss this method briefly. The first step of this procedure is choosing a proper wavelet ***ϕ*** depending on the characteristics of the singularities (e.g. isotropy or the biggest singularity exponent) in the data *f*. The wavelet used in this method works as a lens for detection of singularities which is blind to singularities that have an exponent larger than the order of the wavelet *n*_*ϕ*_. The order of a wavelet ***ϕ*** is defined as the number of vanishing moments of its derivative ***ψ*** whose components are defined as

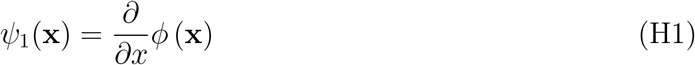

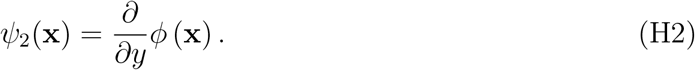

In this study, we determine the range of singularities from the singularity spectrum calculated via MFDFA. Therefore, we can choose the wavelet accordingly. Next, we perform the wavelet transform as following:

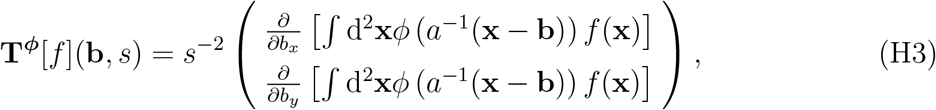

where *s* is the scale, **x** is the position vector, and **b** is its conjugate. We use the modulus of this transform *ℳ*^***ϕ***^[*f*] as a measure to find the singularities in a fashion similar to the aforementioned example. Fig. 11a show the components of the transform and its modulus as well as the argument for the simple example of Fig. 1A. Here, we used Gaussian wavelet which is a first-order wavelet ideal for detecting isotropic singularities with exponents smaller than 1.

**FIG. 11:**
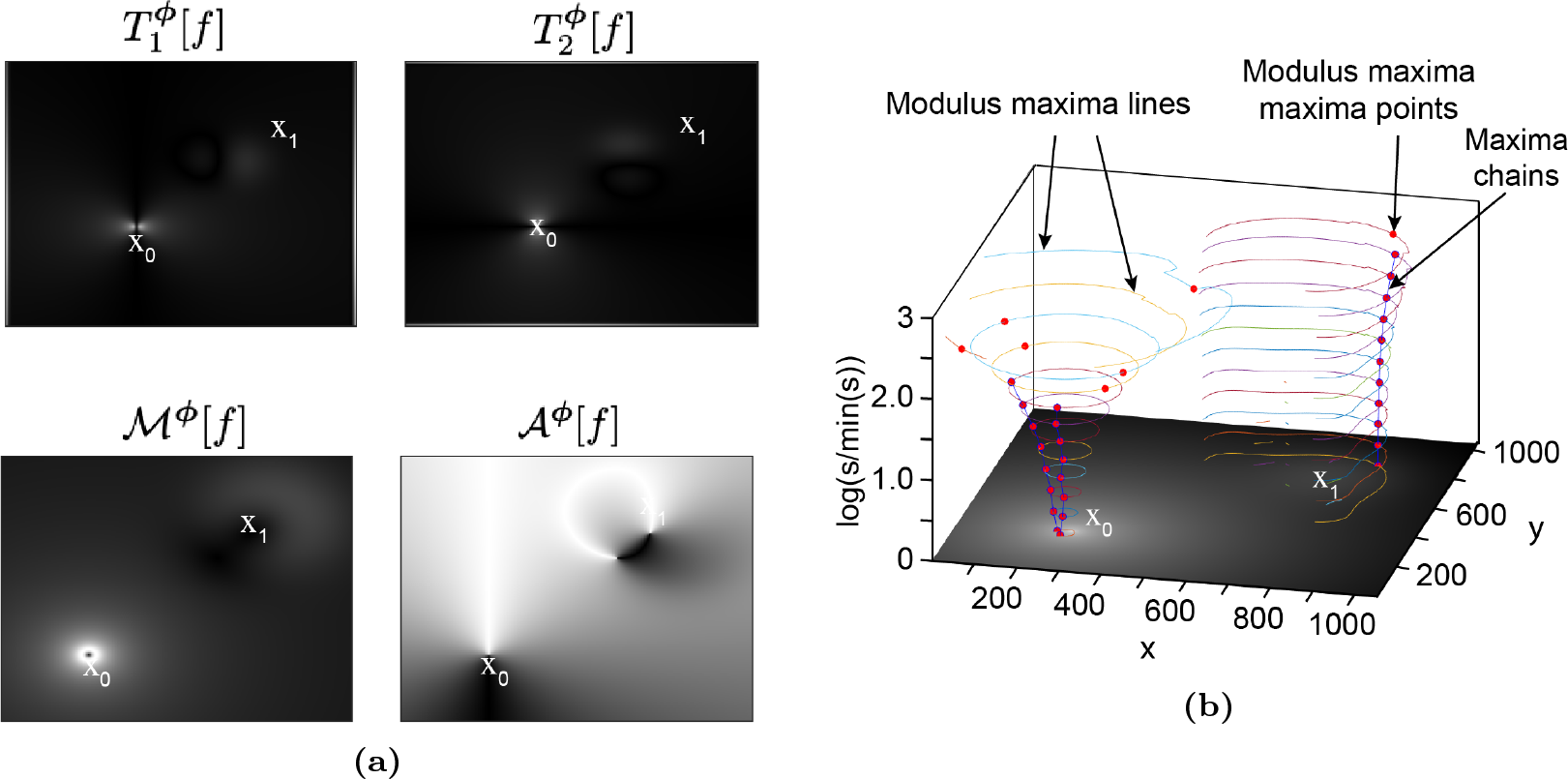
Simple singularity tracking procedure via WTMMMM. **(A)** The components 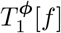 and 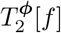 and modulus *ℳ*^***ϕ***^[*f*] and argument *𝒜*^***ϕ***^[*f*] of the wavelet transform of the simple example in Fig. 2A using Gaussian wavelet with *s* = 20 **(B)** The Wavelet modulus maxima lines, modulus maxima maxima points (red circles) and maxima chains through different scales.

After performing the transform, one can find the local maxima of modulus along the argument in order to determine the location of highest intensity changes, similar to the Canny’s multi-scale edge detection [35]. This enables us to construct the maxima lines at each scale which is shown in Fig. 11b for the case of the isolated singularity of Fig. 1A. In this case, at each scale, there are two maxima lines: one is a closed loop around the singular point and the other one which is an open curve around the Gaussian maximum.

The next step is finding the local maxima along the maxima lines. In Fig. 11b, these are shown by red circles. Then one needs to chain these maxima points together throughout different scales to construct the maxima chains and the *skeleton* of the data which is the collection of all maxima chains. The blue lines in Fig. 11b show these chains for our simple example in which two of the lines point towards the singularity point while the other one is pointing towards the steepest side of Gaussian local maximum (the side opposite to the singularity). The last step of singularity detection via WTMMMM is determining whether a chain corresponds to a singularity or not. To do this, one can simply investigate the power-law exponent of the modulus along the maxima chains. The exponent of the power-law is dictated by the order of the wavelet if the chain corresponds to a point with more derivatives than the order of the wavelet. Obviously, this is the case when the point that the chain is pointing toward is a local maximum with infinite derivatives, but it is also the case when the order of the wavelet is less than the exponent of the singularity. Therefore, these two cases are not differentiatable via this method if the wavelet is not chosen properly. Nevertheless, if the point is a singularity with an exponent smaller than the wavelet order, the exponent of the power-law is equal to that of the singularity. This is demonstrated by using our simple example in Fig. 1E where the the blue circles show the modulus along the chains that are pointing to the singularity point and follow a power-law with exponent 0.3. The red triangles however correspond to the chain of the Gaussian maxima that show a power-law with exponent 1 which is unsurprisingly equal to the order of the Gaussian wavelet. It should be noted that as a noisy data gets smoothed by wavelets with larger scale *s*, more information about the smaller scale fluctuations is lost. Therefore, at smaller scales there exist more chains that vanish as one goes to the larger scales.

So far we have discussed the singularity detection aspect of WTMMMM. However, this method with a few extra steps can be used to characterize multifractal features of images as discussed in App. H. Since this method is less accurate than MFDFA [22], we only use MFDFA for determining multifractal features and WTMMM for detecting singularities.

Having constructed the skeleton of the data as shown in Sec. H 1, one can calculate the partition function defined as

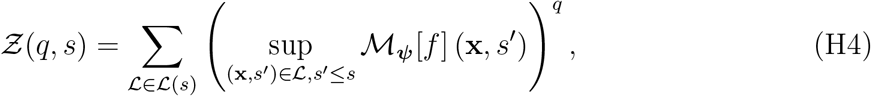

where *ℒ* (*s*) is the set of all maxima chains that exist at scale *s* and persist at scales *s*^*′*^ *< s*. This partition function then can be used to determine the multi-fractal features of the data.

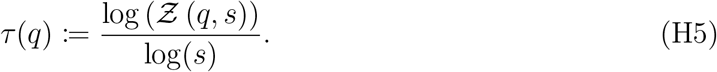

One can define the exponent *τ* (*q*) as

Here, *τ* (*q*) is linear if the data is mono-fractal, and nonlinear if it is multi-fractal. Alternatively, one can use a Legendre transform to obtain the singularity spectrum as following:

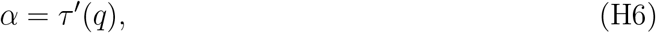

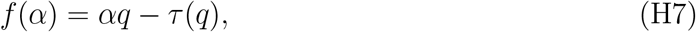

where results in a delta function for a mono-fractal signal and in a wide spectrum for a multi-fractal signal.

In order to show the next final steps of the WTMMMM on a data set with known fractal features, we apply it to a two-dimensional Brownian motion field with *H* = 0.6 and size 1024 *×* 1024. Note that this is a monofractal field with constant generalized exponent i.e. *h*(*q*) = *H* = 0.6. Therefore, according to Eq. 1, there’s only one type of sibgularities in the data with exponent *α* = *H* = 0.6 and the Hausdof dimension *f* (*α* = 0.6) = 2. Accordingly, we can here again use the Gaussian wavelet since its order is larger than the largest singularity exponent in the field. Fig. 12a shows the skeleton that are pointing toward the singularity points as *s →* 0^+^. Fig. 12b depicts the partition function in logarithmic scale showing power law at small scales. Finally, Fig. 12c and Fig. 12d show the comparison between the results of MFDFA, WTMMMM and the theoretical values. The exponent can be written in terms of the generalized Hurst exponent *h*(*q*) as *τ* (*q*) = *qh*(*q*) *− D* which for our case reads *τ* (*q*) = 0.6*q −* 2 which is shown by black line in Fig. 12c. As one can see here, for both measures, MFDFA gives more accurate results. This is again consistent with other studies such as Ref. [22]. However, MFDFA does not provide any information about the position of the singularities while in WTMMMM, one can simply extrapolate the maxima chains to *s* = 0 and find the approximate location of the singularities along with their exponents (by determining the exponent of the power law of the modulus *M*^***ϕ***^[*f*] vs. *s*). It should be noted that some maxima chains exist because of local maxima in the data, not singular points.

**FIG. 12:**
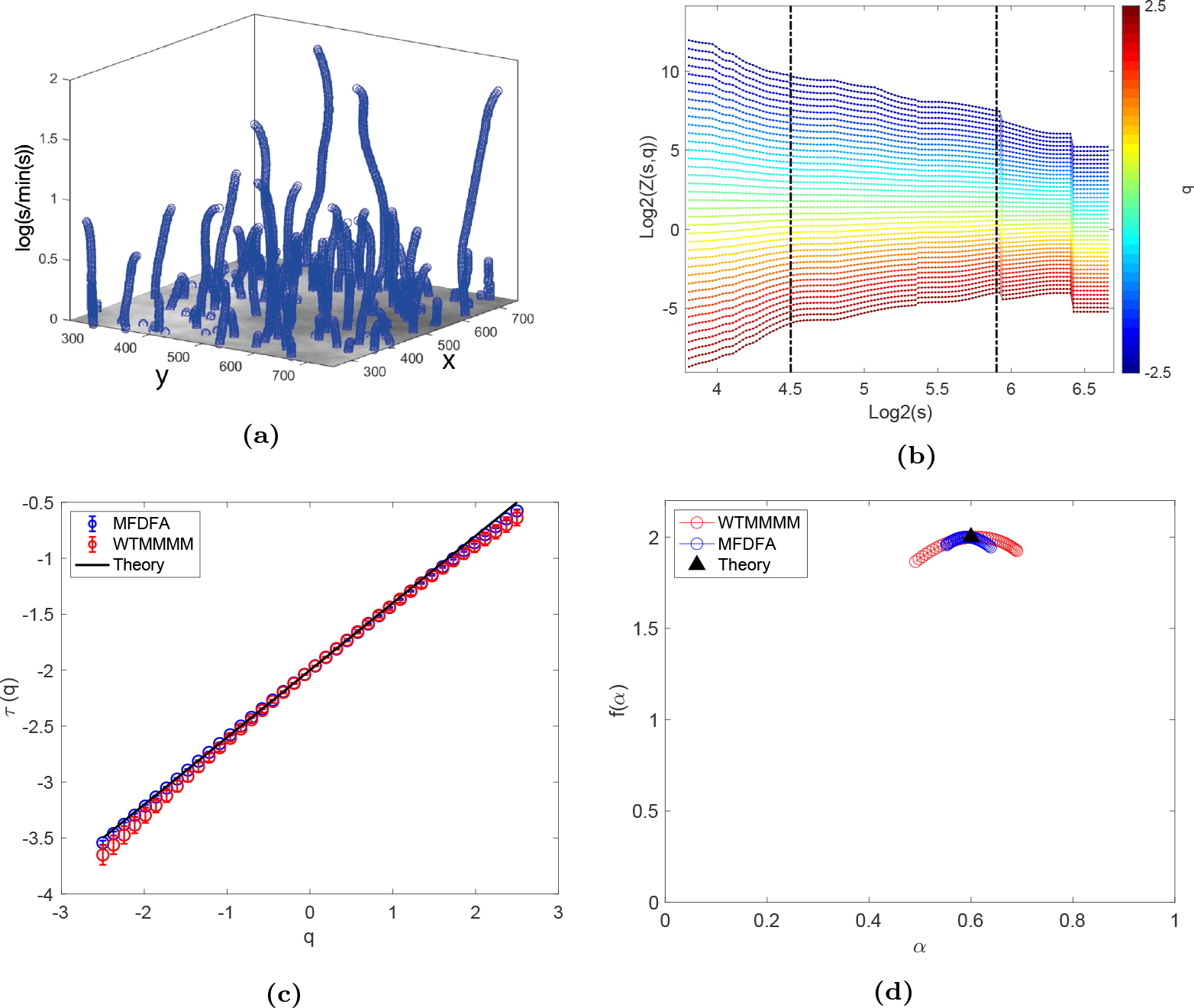
Application of WTMMMM on Fractional Brownian motion field with *H* = 0.6. **(A)** A section of the skeleton of the data. **(B)** The partition function *Ƶ*(*q, s*) in log-log scale. Here, the dashed black lines show the range in which the power laws are fitted to the curves. **(C)** Comparison of the exponent *τ* (*q*) obtained via WTMMMM, MFDFA and its theory value. **(D)** Comparison of the singularity spectrum obtained via WTMMMM, MFDFA and its theory value. Both analyses have good agreement with the theoretical line, but MFDFA performs slightly better.

In these cases, the exponents of the power-laws are determined by the order of the chosen wavelet and, it is easy to distinguish them.

## References

[1] A. Koch and H. Meinhardt, Reviews of modern physics 66, 1481 (1994).

[2] A. Gierer and H. Meinhardt, Kybernetik 12, 30 (1972).

[3] A. Deutsch and S. Dormann, Mathematical modeling of biological pattern formation (Springer, 2005).

[4] T.-H. Nguyen, A. Eichmann, F. Le Noble, and V. Fleury, Physical Review E 73, 061907 (2006).

[5] A. Roth-Nebelsick, D. Uhl, V. Mosbrugger, and H. Kerp, Annals of Botany 87, 553 (2001).

[6] M. Zamir, The Journal of general physiology 67, 213 (1976).

[7] H. P. Schwarcz, D. Abueidda, and I. Jasiuk, Frontiers in Physics 5, 39 (2017).

[8] Y. Liu, Y.-K. Kim, L. Dai, N. Li, S. O. Khan, D. H. Pashley, and F. R. Tay, Biomaterials 32, 1291 (2011).

[9] W. Landis, M. Song, A. Leith, L. McEwen, and B. McEwen, Journal of structural biology 110, 39 (1993).

[10] E. A. Zimmermann and R. O. Ritchie, Advanced healthcare materials 4, 1287 (2015).

[11] A. Lotsari, A. K. Rajasekharan, M. Halvarsson, and M. Andersson, Nature communications 9, 4170 (2018).

[12] Y. Li and C. Aparicio, PloS one 8, e76782 (2013).

[13] C. Cunningham, L. Scheuer, and S. Black, “Bone development:,” (2016) pp. 19–35.

[14] Y. Dang, J. Lattner, A. Lahola-Chomiak, D. Alves-Afonso, E. Ulbricht, A. V. Taubenberger, S. Rulands, and J. Tabler, bioRxiv, 2023 (2023).

[15] C.-L. Benhamou, S. Poupon, E. Lespessailles, S. Loiseau, R. Jennane, V. Siroux, W. Ohley, and L. Pothuaud, Journal of bone and mineral research 16, 697 (2001).

[16] C. B. Caldwell, E. L. Moran, and E. R. Bogoch, Journal of bone and mineral research 13, 978 (1998).

[17] N. Reznikov, M. Bilton, L. Lari, M. M. Stevens, and R. Kröger, Science 360, eaao2189 (2018).

[18] J. W. Kantelhardt, S. A. Zschiegner, E. Koscielny-Bunde, S. Havlin, A. Bunde, and H. E. Stanley, Physica A: Statistical Mechanics and its Applications 316, 87 (2002).

[19] Z. Fayyaz, M. Bahadorian, J. Doostmohammadi, V. Davoodnia, S. Khodadadian, and R. Lashgari, Journal of neuroscience methods 312, 84 (2019).

[20] A. H. Burstein, J. Zika, K. Heiple, and L. Klein, JBJS 57, 956 (1975).

[21] Z. Wang, P. Ustriyana, K. Chen, W. Zhao, Z. Xu, and N. Sahai, ACS Biomaterials Science & Engineering 6, 4247 (2020).

[22] P. Oświecimka, J. Kwapień, and S. Drożdż, Physical Review E 74, 016103 (2006).

[23] G.-F. Gu and W.-X. Zhou, Physical Review E 74, 061104 (2006).

[24] A. Madanchi, M. Absalan, G. Lohmann, M. Anvari, and M. R. R. Tabar, Solar Energy 144, 1 (2017).

[25] S. Mallat and W. L. Hwang, IEEE transactions on information theory 38, 617 (1992).

[26] W. Landis, Bone 16, 533 (1995).

[27] K. R. Wilmarth and J. R. Froines, Journal of Toxicology and Environmental Health, Part A Current Issues 37, 411 (1992).

[28] D. Alves-Afonso, A. Q. Ryan, A. Lahola-Chomiak, M. Prakash, F. Jug, C. D. Modes, and J. M. Tabler, bioRxiv, 2021 (2021).

[29] C. Choquet, L. Boulgakoff, R. G. Kelly, and L. Miquerol, Journal of Cardiovascular Development and Disease 8, 95 (2021).

[30] H. Morales-Navarrete, F. Segovia-Miranda, P. Klukowski, K. Meyer, H. Nonaka, G. Marsico, M. Chernykh, A. Kalaidzidis, M. Zerial, and Y. Kalaidzidis, Elife 4, e11214 (2015).

[31] G. Piszter, K. Kertész, Z. Vértesy, Z. Bálint, and L. P. Biró, Analytical Methods 3, 78 (2011).

[32] M. S. Movahed, G. Jafari, F. Ghasemi, S. Rahvar, and M. R. R. Tabar, Journal of Statistical Mechanics: Theory and Experiment 2006, P02003 (2006).

[33] W. J. Landis, K. J. Hodgens, M. J. Song, J. Arena, S. Kiyonaga, M. Marko, C. Owen, and B. F. McEwen, Journal of structural biology 117, 24 (1996).

[34] A. Arneodo, N. Decoster, and S. . Roux, The European Physical Journal B-Condensed Matter and Complex Systems 15, 567 (2000).

[35] J. Canny, IEEE Transactions on pattern analysis and machine intelligence, 679 (1986).

